# Structural inventory of cotranslational protein folding by the eukaryotic RAC complex

**DOI:** 10.1101/2022.06.24.497458

**Authors:** Miglė Kišonaitė, Klemens Wild, Karine Lapouge, Genís Valentín Gesé, Nikola Kellner, Ed Hurt, Irmgard Sinning

**Author notes:** Correspondence: Prof. Dr. Irmgard Sinning, Biochemiezentrum der Universität Heidelberg (BZH), Im Neuenheimer Feld 328, D-69120 Heidelberg, Germany.

## Abstract

Folding of nascent chains emerging from the ribosome is a challenge in cellular protein homeostasis, which in eukaryotes is met by an Hsp70 chaperone triad directly binding at the ribosomal tunnel exit. The conserved ribosome-associated complex (RAC) consists of the non-canonical Hsp70 Ssz1 and the J-domain protein Zuotin (Zuo1), which in fungi acts together with the canonical Hsp70 protein Ssb. Here, we determined high-resolution cryo-electron microscopy structures of RAC bound to the 80S ribosome. RAC adopts two distinct conformations accommodating continuous ribosomal rotation by a flexible lever arm. The heterodimer is held together by a tight interaction between the Ssz1 substrate-binding domain (SBD) and the N-terminus of Zuo1, with additional contacts between the Ssz1 nucleotide-binding domain (NBD) and the Zuo1 J- and ZHD domains that form a rigid unit. The Zuo1 HPD-motif conserved in J-proteins is masked by the Ssz1 NBD, different from the canonical Hsp70 J-protein contact, however, allowing to position Ssb for activation by Zuo1. Our data provide the basis for understanding how RAC cooperates with Ssb at the ribosome in dynamic nascent chain interaction and protein folding.

## Introduction

Efficient protein folding is a challenge for proteostasis in all organisms, which already during translation is ensured by ribosome-associated chaperones that modulate protein synthesis and are among the first contacts of the emerging polypeptides^1,2^. RAC is conserved in eukaryotes, and in *S. cerevisiae* comprises a stable heterodimer formed by the non-canonical Hsp70 homolog Ssz1 and the J-domain protein (JDP) Zuo1^3,4^. Ssz1 differs from canonical Hps70s in several ways: it binds ATP but does not hydrolyze, and ATP binding is not required for its function^5^; it has a unique domain arrangement and a truncated substrate binding domain (SBD) with only a rudimentary β-sandwich domain (SBDβ); it lacks the α-helical lid domain (SBDa) and the conserved linker^3,6^, which is central to the allosteric regulation of canonical Hsp70 activity^7^. Instead, the linker in Ssz1 is extended and adopts an αβ-structure that intertwines with the Zuo1 N-terminus, which complements SBDβ and moulds this unusual Hsp70/JDP pair into a stable, functional unit^6,8^ (**Fig. 1a**). Zuo1 is a class C JDP and the only Hsp40 that activates the ribosome-associated Hsp70 protein Ssb (encoded by two isoforms *SSB1* and *SSB2,* that are nearly identical)^3,9^. In general, JDPs play a central role in specifying and directing Hsp70 functions^10–12^. They comprise a universally conserved HPD-motif, which is essential for stimulating the ATPase activity in all JDP/Hsp70 pairs^13^. However, the co-chaperone function of Zuo1 requires the presence of Ssz1 ^4^, underlining that RAC and Ssb form a functional chaperone triad at the ribosome^14,15^. Nascent chain (NC) binding by Ssb requires the presence of RAC^16^ and accelerates translation^17^. Both RAC proteins contact the NCs and form a relay system that transfers polypeptides from Zuo1 via Ssz1 to Ssb^8^. The majority of nascent proteins interact with Ssb by multiple binding-release cycles^18^. RAC binding to the ribosome has been thoroughly studied by cross-linking experiments and low-resolution cryo-EM structures showing flexible conformations on idle 80S ribosomes and interactions with both the 40S and 60S subunits^5,19,24^. However, the integration of Ssb, its ATPase cycle, NC and ribosome interactions into the workings of RAC has remained incomplete. A recent *in vivo* cross-linking study suggests a pathway of Ssb movement at the ribosome and places Ssb next to the Ssz1 NBD^24^. However, all these data did not provide a complete picture of the RAC/Ssb triad at the ribosome.

**Fig. 1.**
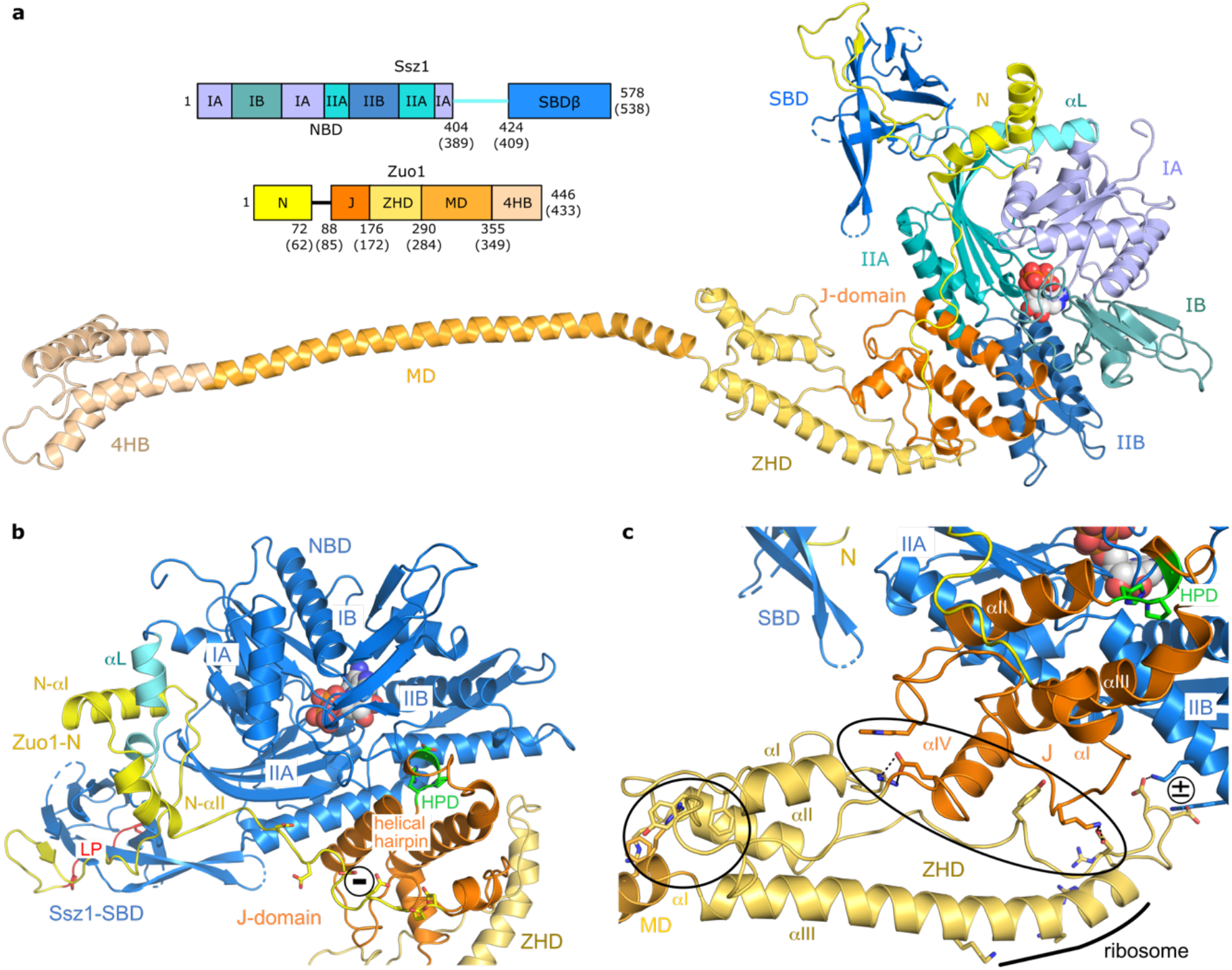
Architecture of full-length RAC reveals new contacts between Ssz1 and Zuo1. **a,** Cryo-EM structure of *Chaetomium thermophilum* RAC in ribbon representation and its domain architecture (residue numbers are given for *C. thermophilum;* corresponding residues in *S. cerevisiae* are in brackets). For purpose of representation, only RAC-1 conformation is shown. Ssz1 comprises a nucleotide binding domain (NBD; shades of blue), a linker (aL; cyan), and a substrate binding domain β (SBDβ; dark blue). NBD lobes IA, IIA, IB and IIB are shown in different shades of blue. Zuo1 comprises an N-terminal domain (N; yellow), J-domain (J; orange), Zuo1 homology domain (ZHD; pale yellow), middle domain (MD; pale orange), and four-helix bundle (4HB; tan). Disordered residues are indicated as dotted lines. ATP is shown in sphere representation. **b,** The Ssz1-Zuo1N interface is enlarged by an extension of Zuo1N-aI and Zuo1N-αII. Zuo1-J shows the canonical J-domain fold with a central helical hairpin and contacts Ssz1-NBD. It bridges lobes IIA to IIB and contains the conserved HPD-motif (CtZuo1 His133-Pro134-Asp135; green). The HPD-motif breaks the first helix at its C-terminus and is completely masked by its Ssz1-NBD interaction. LP-motif binding to the Ssz1-SBD is highlighted in red. **c,** Zuo1-J and -ZHD are directly linked and form a rigid entity. The J-ZHD contact involves salt bridges and stacking aromates (large black ellipse). An additional small contact (present in RAC-1 conformation only) between Zuo1-ZHD and Ssz1-NBD involves two aspartates adjacent to ZHD-αIII (annotated by +/-). The Zuo1 ZHD-MD contact is indicated by a small black circle.

## Results

We now determined high-resolution structures of RAC bound to translating 80S ribosomes using native *Chaetomium thermophilum* (*C. thermophilum, Ct*) complexes^25^ pulled-out on Ssz1 for subsequent cryo-EM structure determination at 3.2 and 3.3 Å resolution (**Fig. 1, Extended Data Figure 1** and **Extended Data Table 1**). We obtained multiple 80S-RAC structures (with different ribosomal rotation states and RAC conformations) including mixtures of nascent chains visible from the peptidyl-transferase center (PTC) to the very tunnel exit, and with extra-ribosomal factor RACK1 and the protective factor Stm1 bound as recently described for *Ct*80S ribosomes^26^. The quality of the cryo-EM map representing RAC allowed us to build a complete model of this multidomain complex (**Fig. 1a**). Our recent X-ray-structures of the RAC core comprising Ssz1 with its SBD completed by the Zuo1 N-terminus, could be readily placed as rigid-bodies^6,8^. Although for Zuo1-ZHD (Zuotin homology domain) and the C-terminal four helix-bundle (4HB) structural models were available^20,22,27^, large parts of Zuo1 including the J-domain, the MD and linkers between domains had to be built *de novo* (**Fig. 1b, c** and **Extended Data Fig. 2**).

### Complete RAC reveals contacts between Ssz1-NBD and Zuo1 J-ZHD

In our RAC structures, Zuo1 contacts the Ssz1-SBD mainly by the previously described tight interaction with Zuo1N (residues 1 to 72; 3060 Å^2^ interface area with a ΔG of −42.4 kcal/mol, 80% of the total Ssz1-Zuo1 interface)^6,8^. A conserved polyproline type-II helix (LP-motif) at the Zuo1 N-terminus binds to the Ssz1-SBD as a pseudo-substrate^6,8^. The Ssz1-Zuo1N interface is now enlarged by an extension of Zuo1N-aI and an additional α-helix (Zuo1N-αII, residues 62 to 72; **Fig. 1** and **Extended Data Fig. 2**) that grab the Ssz1 specific linker helix (αL) connecting NBD and SBD. A linker between Zuo1N and the J-domain (residues 73 to 87) is highly negatively charged and barely contacts the Ssz1-SBD and Zuo1 J-domain (residues 88 to 175). The J-domain shows the canonical fold of JDPs with a central helical hairpin^28^ that forms the only contact between the J-domain and Ssz1-NBD (685 Å^2^, ΔG of −2.7 kcal/mol). This hairpin bridges lobe IIA to IIB and contains the conserved HPD-motif (*CtZuo1* His133-Pro134-Asp135), which is crucial for Hsp70 activation^28^. The HPD-motif breaks the first helix at its C-terminus and is completely masked by its Ssz1-NBD interaction (**Extended Data Fig. 3a;** see below). This contact differs from the classical Hsp70/JDP activating complex^29^, where the HPD-motif interacts with the conserved Hsp70 linker region, inserts the helical hairpin between NBD lobes IA and IB, and contacts also SBDβ (**Extended Data Fig. 3b)**. Of note, this canonical contact is also small and unstable (925 Å^2^, ΔG of −1.3 kcal/mol), which is reflected by a generally transient Hsp70/JDP interaction^7^. However, in both cases the contact involves mostly polar or ionic residues, and is centred around the HPD-motif (and a following positive residue) with the aspartate forming a salt bridge.

In contrast to the Zuo1 N-J connection, the Zuo1-ZHD (residues 176 to 289) is directly linked to the J-domain, which together form a rigid entity **(Fig. 1a-c).** The ZHD closely corresponds to an X-ray structure for yeast Zuo1-ZHD (root mean squared deviation of 2.0 Å)^22^ and comprises a three-helix bundle with an extended C-terminal α-helix (ZHD-αIII). The ZHD was previously characterized as ribosome-binding domain^22^, but its interactions within RAC were not resolved. Our structures reveal an intimate contact with Zuo1-J, mostly to J-αIII flanked by loop interactions involving salt bridges and stacking of aromatic residues (buried surface area 623 Å^2^, ΔG of −5,2 kcal/mol). We also observe an additional small contact between Zuo1-ZHD and Ssz1-NBD, which involves two aspartates adjacent to ZHD-αIII (**Fig. 1c**). This contact changes between the two distinct conformations of RAC on the 80S ribosome (see below). Between the ZHD and the following helical middle domain (MD, residues 290 to 354) a tight π-cation stacking network is observed fixing the first two turns of MD-aI to the ZHD (**Fig. 1c**). The MD connects the three N-terminal Zuo1 domains to the rigid C-terminal four-helix bundle (4HB; residues 355 to 446), which anchors RAC on the 40S subunit by interacting with the rRNA expansion segment ES12. Taken together, our data allow to build a complete model of RAC with precisely defined domain boundaries and to describe interactions within Zuo1 as well as with Ssz1, which are different from canonical Hsp70/JDP interactions.

### Two distinct conformations of RAC on the 80S ribosome

Consistent with previous data^3,6,20,21^, our RAC-80S complexes display an extended RAC structure that spans more than 200 Å and contacts both ribosomal subunits. RAC adopts two distinct conformations (denoted RAC-1 and RAC-2) on a rotating ribosome (**Fig. 2, main panels** and **Extended Data Fig. 4**). This ratchet-like motion is a conserved feature of all ribosomes and is intrinsic to mRNA/tRNA translocation^30^. 3D variability analysis^31^ allowed us to visualize continuous movement of the 40S subunit in respect to 60S for both RAC conformations (**Extended Data Fig. 5a** and **Extended Data Movie 1**). It was previously thought that RAC stabilizes the 80S ribosome in the non-rotated state and that its movement is coupled to ribosomal rotation^21^. However, our structures demonstrate that idle 80S ribosomes containing RAC in either conformation exhibit the same distribution of rotational states (**Extended Data Fig. 5b**). The rotation of the entire 40S body, except the ES12 movements, in both cases reaches to about 7° and the swiveling of the 40S head reaches up to 18°. For better comparison, RAC-1 and RAC-2 were built on the non-rotated ribosome.

**Fig. 2.**
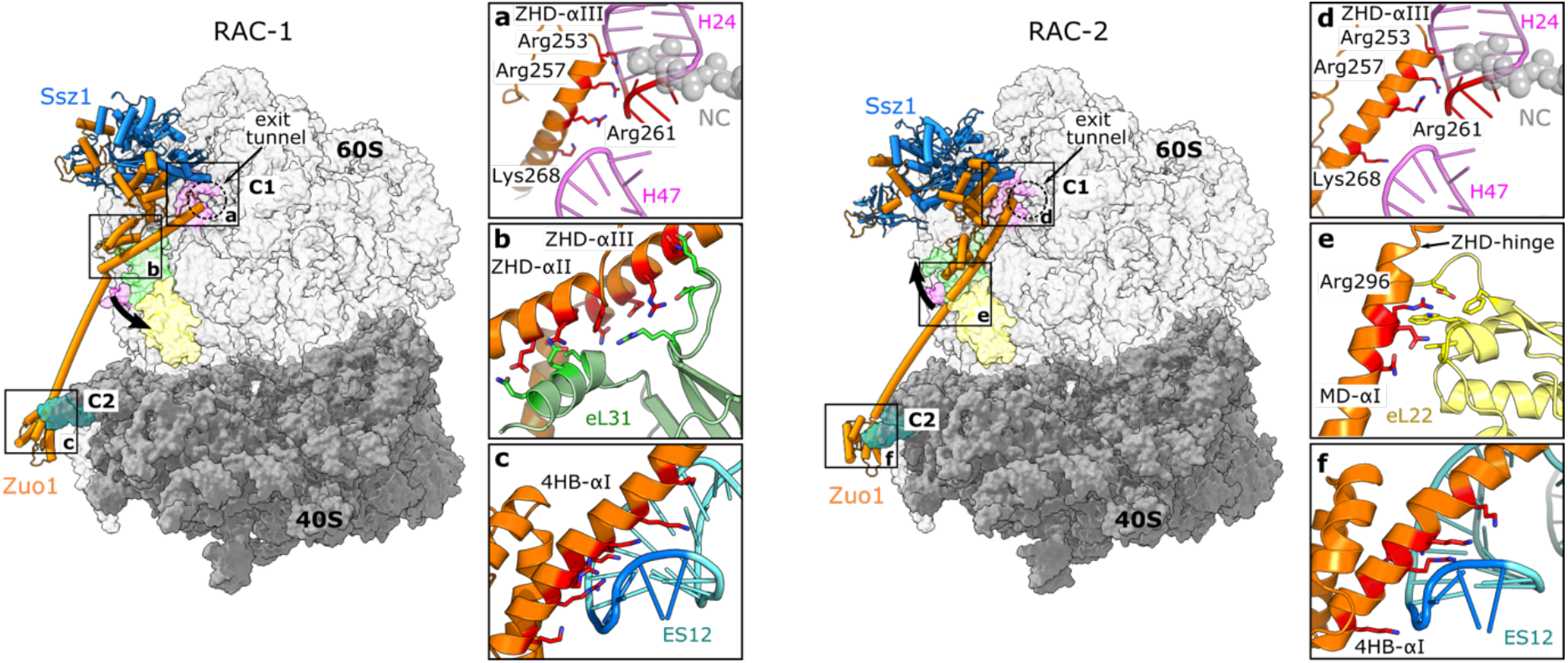
RAC interactions with the 80S ribosome. Cryo-EM structures of *Ct*RAC bound to the 80S ribosome in two distinct conformations – RAC-1 (left) and RAC-2 (right). The main 80S contacts are highlighted with squares that correspond to the zoom images **a** to **f**. **a, d,** ZHD-80S interaction (C1 contact) with H24 and H47 of the 26S rRNA in RAC-1 (**a**) and RAC-2 (**d**). C1 is formed by the N-terminal end of the lever arm (ZHD-αIII) at the rim of the ribosomal tunnel exit. **b,** ZHD interaction with the eL31 ribosomal protein in RAC-1. **e,** ZHD-MD interaction with the eL22 ribosomal protein in RAC-2. **c, f,** 4HB interaction (C2 contact) with ES12 of the 18S rRNA in RAC-1 (**c**) and RAC-2 (**f**). C2 is formed at the C-terminal end of the lever arm (Zuo1-4HB) and the closing tetraloop (1695-GCAA) of ES12 in the 40S subunit.

In both RAC conformations, interactions with the ribosome are exclusively formed through Zuo1 by a lever arm (residues 253 to 371) that we define based on our structures to include ZHD-αIII (residues 253 to 289), the entire MD (residues 290 to 354), and 4HB-αI (residues 355 to 371). Ssz1 does not interact with the ribosome, but is kept in close proximity to the ribosomal tunnel exit by its interaction with Zuo1N^6,8^ and by the two small contacts between Ssz1-NBD with the Zuo1 J-ZHD unit.

While a previous study suggested that the RAC-ribosome interaction changes with ribosomal rotation^21^, our data clearly show that the Zuo1 lever arm anchors RAC at the ribosome with two main contacts (C1 and C2) that are maintained in both RAC conformations independent of the ribosomal rotation state. C1 is formed by the N-terminal end of the lever arm at the rim of the ribosomal tunnel exit (**Fig. 2a, d**) with three conserved arginines from ZHD-αIII (Arg253, 257, and 261; for homology see **Extended Data Fig. 2**). These arginines form a so-called ARM (arginine-rich motif)^32^ that affixes Zuo1 in the major groove of the tetranucleotide loop (tetraloop, 376-GAAA) at the tip of helix H24 of 26S rRNA. The interaction is completed by the positive N-terminal helix dipole of ZHD-αIII, which positions the helix on the phosphoribose backbone. In yeast, the corresponding arginines 247 and 251 also contact H24 of the 26S rRNA^22^, and disruption of this contact completely abolishes RAC binding to the ribosome in yeast, both *in vitro* and *in vivo*^33^.

C2 is formed at the C-terminal end of the lever arm between Zuo1-4HB and the closing tetraloop (1695-GCAA) of 18S rRNA ES12 in the 40S subunit (**Fig. 2c, f**). Similar to C1, C2 also involves an elaborate ARM interaction between 4HB-aI and ES12. The helix contributes two arginines (Arg362, 365) and five lysines (Lys350, 354, 358, 359, and 369) to this interaction. While ES12 shortening severely affected translation fidelity and readthrough effects of stop codons, the RAC-ribosome interaction was only mildly destabilized^22^.

The C1 and C2 contacts appear invariant in both RAC conformations. However, the lever arm undergoes a complex motion, which can be described by a bending elbow located in the MD (here denoted as MD-elbow at Lys305; **Extended Data Fig. 6a**). While in RAC-1 the MD-elbow is bent by 37°, it is straightened up in RAC-2 (**Fig. 2**, main panels). In addition, two minor hinges (<20°) localize at both ends of the lever arm, between ZHD and MD (ZHD-hinge at Glu290) and between MD and 4HB (4HB-hinge at Asn355) (**Extended Data Fig. 6b, c**). Interestingly, when RAC-1 and RAC-2 are superposed on Ssz1 (**Extended Data Fig. 6**), Ssz1 and Zuo1 J-ZHD (as well as the 4HB by itself) overall behave as rigid bodies (root mean squared deviations <1.3 Å). However, as both ends of Zuo1 are fixed on the ribosome, the invariant C1 and C2 contacts must somehow accommodate changes within the MD-elbow. Indeed, when comparing the RAC-1 and RAC-2 contacts with the ribosome, the Zuo1-ZHD rotates around C1 (residues 246-261) in respect to the J-ZHD unit (45° rotation at borders) (**Extended Data Fig. 6d**), while C2 is maintained by a significant bending of ES12 (**Fig. 2**, main panels; and see below).

Apart from C1 and C2, there are several interactions between the lever arm and the ribosome that are adjusted. In RAC-1, Zuo1-ZHD interacts with protein eL31 via a mixed polar-apolar helical bundle (ZHD-aII and eL31 N-terminal helix) and multiple salt bridges between the lever arm (ZHD-αIII) and an internal eL31 loop (**Fig. 2b**). This interaction nicely correlates with previously observed cross-link data^22^. Interestingly, the eL31 N-terminal helix is rotated by 50° towards the ZHD compared to RAC-2 (and the idle 80S ribosome^26^) (**Extended Data Fig. 6e**). Furthermore, the MD-elbow rests on the 26S rRNA 3’-end (H101) with Arg310 seemingly stacking on a bulged-out cytosine (C3324) (**Extended Data Fig. 7a**).

In RAC-2, these interactions have disappeared (**Extended Data Fig. 7c, d**) and straightening the MD-elbow moved the lever arm by up to 40 Å on top of protein eL22, which fixes the ZHD-hinge by two internal loops and its very C-terminus (**Fig. 2e**). In particular, Zuo1 Arg296 is involved in π-cation stacking with a tryptophan and in a salt bridge. Previous cross-linking studies failed to detect the eL22 contact, probably due to technical reasons^22^. Finally, adjacent to C1 a weak contact between H47 and a single lysine (Lys268) is observed, which is lost in RAC-1 (**Fig. 2a, d**). Another striking difference is observed next to the tunnel exit at the contact between Zuo1-ZHD and Ssz1-NBD (**Fig. 3a, b**). In RAC-1, this contact comprises two salt-bridges (Zuo1-Asp248/Ssz1-Lys255, Asp249/Lys259), which are absent in RAC-2 as Ssz1-NBD has detached from Zuo1-ZHD and moved away from the tunnel exit by 10 Å.

**Fig. 3.**
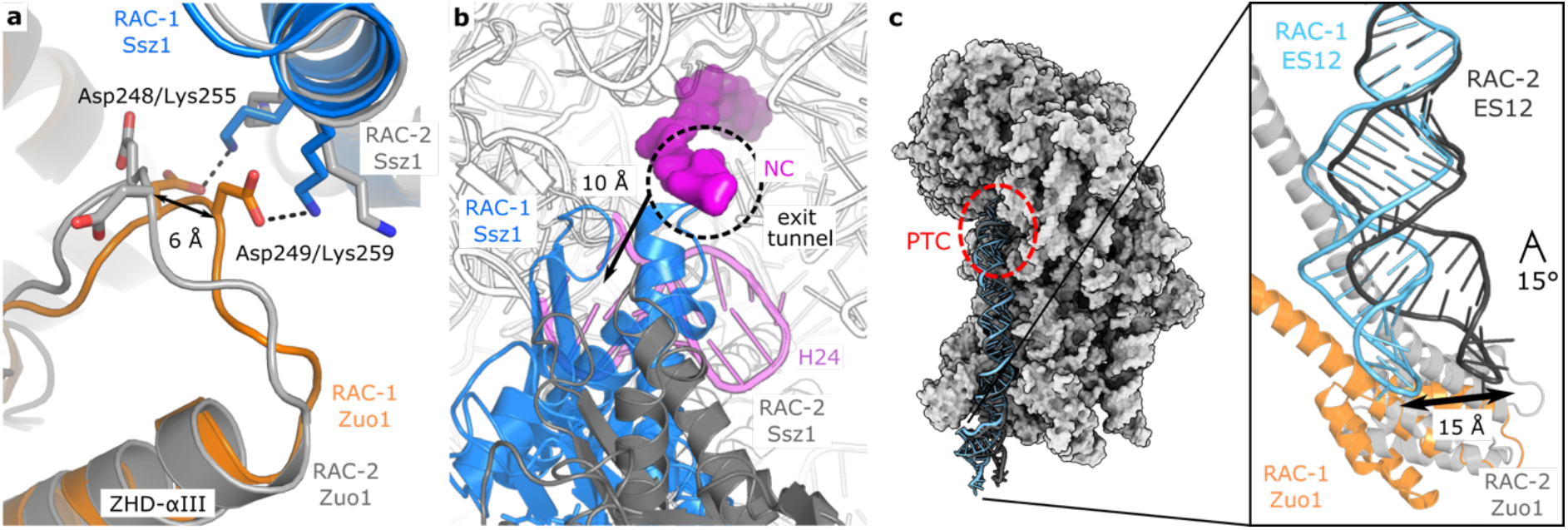
Details of structural differences between RAC-1 and RAC-2. RAC-1 is shown in color (Ssz1 – blue, Zuo1 – orange), while RAC-2 is shown in grey. **a** and **b**, Contact between Zuo1-ZHD and Ssz1-NBD close to the tunnel exit. In RAC-1, this contact comprises two salt-bridges (Zuo1-Asp248/Ssz1-Lys255, Asp249/Lys259), which are absent in RAC-2 (shift of 6 Å, **a**). The contact between Zuo1 and Ssz1 is abolished as Ssz1-NBD has detached from Zuo1-ZHD and moved away from the tunnel exit by 10 Å (**b**). Nascent chain (NC) is shown in magenta and represented as surface. 26S rRNA H24 that is involved in the C1 contact is shown in pink. **c,** The 40S-Zuo1 contact. 40S shown in surface representation (left; grey) with ES12 of the 18S rRNA shown in sticks, and the peptidyl transferase center (PTC) highlighted by a red circle. Zoom in view (right) of the Zuo1-4HB interaction (C2 contact) with ES12 in both RAC conformations. The 4HB-ES12 contact stays invariant, but the tip of ES12 adapts by a 15° bend and moves by 15 Å.

While at C2 the contact with ES12 stays invariant in both RAC conformations and throughout ribosomal rotation, the tip of ES12 adapts by a 15° bend in a movement independent from 40S body rotation (**Fig. 3c**). The tip of ES12 thus moves by 15 Å. Interestingly, next to its flexible tip, ES12 is held in place by another ribosome-internal ARM, this time provided by eL24 of the 60S subunit that is threaded through a widened ES12 major groove and with its long C-terminal helix anchors on the 40S body (**Extended Data Fig. 8**). Furthermore, ES12 forms the end of the long 18S rRNA helix H44 located in between the 40S and 60S subunits (200 Å length) that reaches up to the codon-anticodon base pairs, and contacts Stm1 that occupies the P-site as described recently^26^. H44 is known to ensure the accuracy of translation elongation and termination^22^, however further investigation is needed to delineate the exact role of RAC in translational fidelity. Overall, we observe RAC in two distinct conformations on a rotating ribosome and resolve mechanistic details of RAC-80S interactions.

### Model of Ssb stimulation by Zuo1

RAC forms a functional chaperone triad with Ssb, which needs activation by Zuo1-J for productive interaction with nascent chains. Our structures of RAC at the 80S, and structures of the *E. coli* DnaK/DnaJ complex^29^ and of yeast Ssb (open, ATP-bound state)^34^ allow us to derive a structure-based model of the RAC/Ssb triad at the ribosome. First, the DnaJ J-domain is superposed on Zuo1-J (RAC-2 chosen, RAC-1 also possible), and second, DnaK (in the DnaK/J complex) is replaced by Ssb^34^ to obtain a model for Ssb activation by the Zuo1 HPD-motif (**Extended Data Fig. 9**). In the superposition of the J-domains, the NBDs of DnaK and Ssz1 would clash. The Ssz1-NBD that masks the Zuo1 HPD-motif (described above) needs to detach from the Zuo1 J-ZHD unit, which is anchored at the ribosomal tunnel exit by C1. Noteworthy, in RAC-2 the slight detachment of Ssz1-NBD from Zuo1-ZHD (moved away from the tunnel exit by 10 Å compared with RAC-1) already opens this weak contact and provides access to the tunnel exit. The short Zuo1 N-J linker (13 residues) will keep Ssz1-Zuo1N in close neighborhood. Superimposing Ssb on DnaK places Ssb-SBDβ directly on top of the tunnel exit ready for interaction with short nascent chains consistent with previous cross-link and ribosome profiling data (**Extended Data Fig. 9c**)^3,17^. In the ATP-bound open state, the Ssb-SBDa lid domain is not interfering with any contacts and points away from the ribosome. This seems counterintuitive as the lid domain harbors the key ribosome binding motif of Ssb. However, the structures of Ssb-ATP and DnaK-ATP have been obtained by fixing the domain arrangement by an engineered disulfide bridge^34,35^. In addition, autonomous ribosome binding of Ssb is not required for its function in presence of RAC^36^.

In contrast to most Hsp70 chaperones that can be activated by several JDPs, it has been shown that Zuo1 is the only JDP that activates Ssb and stimulates ATP hydrolysis^5^. However, the basis of this specificity was not clear. Our model with Ssb in the activating position does not show any clashes with Zuo1 or the ribosome, and the Ssb-Zuo1-J interface shows all characteristic interactions described for the DnaK-DnaJ complex^29^ (**Extended Data Fig. 10**). In addition to these canonical Hsp70/JDP interactions, our model also visualizes Ssb-specific interactions with Zuo1. Interestingly, these specific interactions mainly involve a KRR-motif (residues 429-431 in *Sc*Ssb; KKR-motif in *Ct*Ssb) in Ssb-SBDβ that has previously been described as a ribosome attachment point^24,36^. In our model however, the two lysines embrace Zuo1-J Trp98, while the arginine forms a salt bridge with Zuo1-ZHD Asp248 (**Extended Data Fig. 10c**) that replaces the interaction with Ssz1-NBD observed in RAC-1 (but not in RAC-2). Therefore, our structure-based model suggests that the KRR-motif contributes to the specific activation of Ssb by Zuo1.

## Discussion

The RAC-80S structures described here provide the details of RAC architecture and the RAC/80S interaction. The contacts observed between 80S ribosomes and RAC localize this specific Hsp40/Hsp70 activity at the ribosomal tunnel exit and provide an answer to the function of Ssz1 and the specificity of the Zuo1/Ssb pair. Together with previously obtained crystal structures of JDP/Hsp70 complexes^29,37^ and Ssb^34^, the RAC-80S structures allow us to extend on our RAC/Ssb model and propose a mechanism for the action of the RAC-Ssb chaperone triad on the ribosome (**Fig. 4**). The mechanism is based on our observation that the strong ARM contacts of Zuo1 stay invariant during ribosomal rotation and that the ZHD/J-domain entity behaves as rigid body. Thus, it can be assumed that RAC remains attached to the RNC during protein biosynthesis and the J-domain position adapts to the observed RAC-conformations. The second premise is that the activating JDP/Hsp70 interaction, mediated by the HPD-motif, is universally conserved and that the available crystal structures can serve as general template. While in non-activating case of Zuo1/Ssz1, the HPD-motif is completely masked by its Ssz1-NBD interaction. Furthermore, nascent chain (NC) binding contributes to RAC/Ssb interaction at the ribosome, and specific sequence requirements for Ssb/NC interaction were determined by ribosome profiling^17^. Ssb binds to degenerated sequence motifs enriched in positively charged and hydrophobic residues positioned at a distance of 35-53 residues from the PTC^17^, and cross-linking data indicate that the NC is handed over in a relay from Zuo1 via Ssz1 to Ssb^8^. In the absence of functional RAC, Ssb fails to interact with NCs as the high-affinity substrate binding state of Ssb is not induced^15,34^. Our structures now localize the Zuo1-ZHD next to the tunnel exit and show that it not only modulates ribosome and Ssz1 interactions, but also exposes a highly negatively charged surface in a matching distance from the PTC. The adjacent Ssz1-NBD IIB lobe is also negatively charged (and slightly hydrophobic) while the more distal interface to the IA lobe is strongly positively charged. Our current model integrates these observations, and suggests that complementary charges might contribute to NC binding and handover. Positively charged NCs first interact with Zuo1-ZHD, while slightly longer NCs, can bind to adjacent negative and slightly hydrophobic patches in Ssz1-NBD lobe IIB (**Extended Data Fig. 10a**). Further elongation of the NC and the dynamic Zuo1-ZHD/Ssz1-NBD contact, as observed between the RAC-1 and RAC-2 complexes, can then direct the NC towards the positive patch in between Ssz1-NBD lobes IB and IIB, and are probably sufficient to dissociate the weak contact between Ssz1-NBD and the Zuo1 HPD-motif. This would allow Ssb to join in and engage in the canonical activating, transient J-domain contact (**Extended Data Fig. 10b**). Activation of ATP-hydrolysis in Ssb drives efficient NC interaction (Ssb in the ADP state) when dislodging from the ribosomal surface. The J-domain can then again be masked by Ssz1 to avoid unproductive engagements e.g., with another Ssb molecule.

**Fig. 4.**
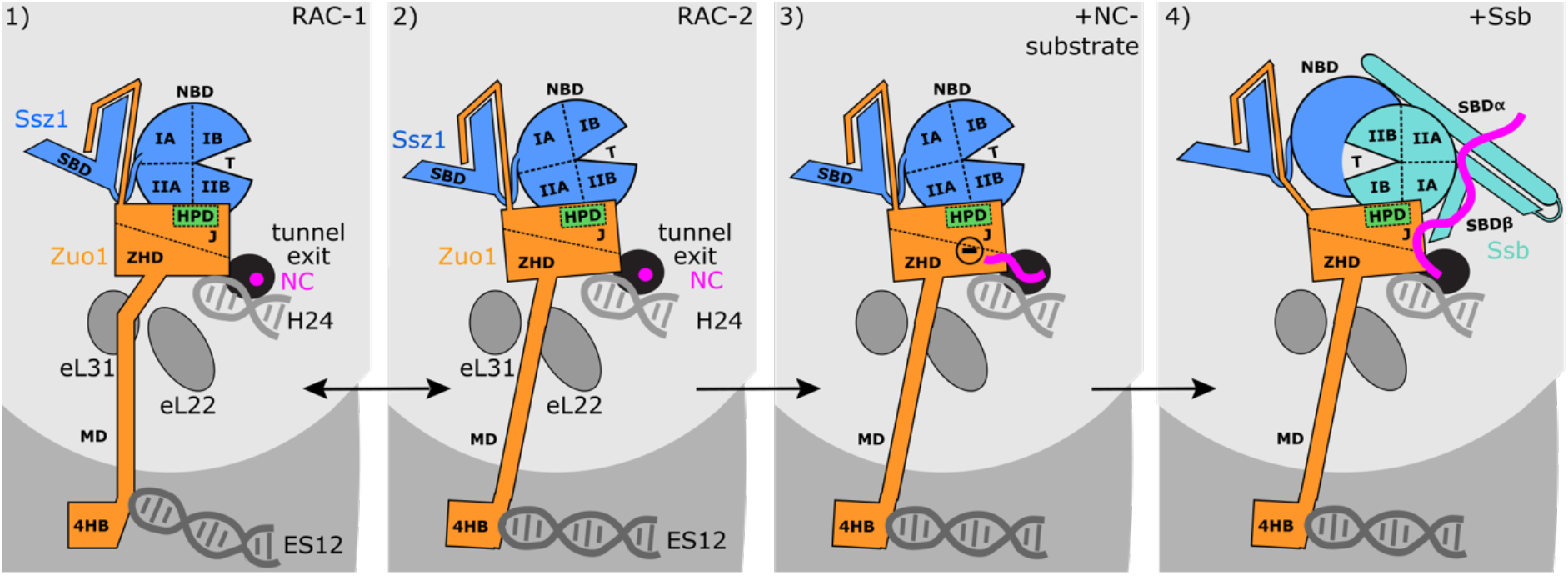
Structure-based model for RAC/Ssb action at the 80S. Integrating our cryo-EM structures with the current data on RAC and Ssb allows to devise a detailed model of RAC/Ssb action at the ribosome. RAC binds to the 80S in two distinct conformations with Zuo1 oscillating between RAC-1 and RAC-2 (panels 1, 2). The HPD-motif of Zuo1-J (green) is masked by Ssz1-NBD. A RAC/Ssb substrate (positively charged NC) emerging from the exit tunnel first interacts with a negatively charged patch in Zuo1 (panel 3). Elongation of the NC allows it to reach a positively charged patch in Ssz1 (not indicated). The NC pushes the Ssz1-NDB away and thereby frees the HPD-motif. This allows for Zuo1-Ssb interaction, the growing NC contacts Ssb, which can now be stimulated by Zuo1-J (panel 4). Ssb is positioned next to the tunnel exit, with its SBD conveniently placed close to the emerging NC and its NBD forming a heterodimer with Ssz1-NBD. When the Ssz1-NBD is displaced from the HPD by NC and Ssb binding, the Ssz1-SBD stays tied-up with Zuo1-N. After stimulation of ATP hydrolysis Ssb can detach from the ribosome and Ssz1-NBD returns to shield the HPD-motif.

The position of Ssb at the ribosome has remained quite puzzling despite several cross-link studies^3,5,19–24^. Recent data place Ssb next to the tunnel exit with different binding modes (with bound ATP or ADP)^24^. Furthermore, these data specify interactions between Ssz1-NBD with both Ssb-NBD and -SBDa, and suggest the formation of an Ssz1-Ssb NBD heterodimer. Such placement of Ssb nicely correlates with our cryo-EM structures and supports our structure-based model (**Fig. 4** and **Extended Data Fig. 10c**). Notably, the proposed heterodimer interaction resembles homodimers observed in crystals of the Hsp70s Ssb^34^ and DnaK^38^, and also the Hsp110 Sse1 ^39^, suggesting that NBD dimer formation might be more common in Hsp70 and Hsp110 chaperones.

The two distinct RAC conformations observed in this study do not correlate with ribosomal rotation. Therefore, the question remains to what triggers RAC-1/RAC-2 oscillation, and how the entire chaperone triad is coupled to translation on one hand and to Ssb ATP-binding and hydrolysis on the other hand. It is tempting to speculate that factors missing in our study might be involved, e.g., the complete mRNA∂tRNA2 module and a steadily growing NC that harbors Ssb-substrate sequences. ES12 dynamics is likely to play an essential role with ES12 also being important for fidelity of translation^22,40^. Different studies already investigated the 4HB interaction with ES12^22,40^, however, so far only with perturbed or truncated systems. While it was previously envisaged that a direct coupling between RAC binding and the ribosome active center occurs via the central rRNA helix H44 including ES12 at its tip^23^, our structures suggest the RAC influence on fidelity to depend on its constant binding probably by modulating the speed of ratcheting. Further functional and especially high-resolution structural studies of all components of stalled on-pathway complexes are needed to finally unveil the complete movie of this unique co-translational chaperone triad in protein biosynthesis. The absence of a Ssz1 homolog in humans and the presence of additional domains in *hs*Zuo1 together with off-ribosomal transcriptional functions of Zuo1 and Ssz1^23,41^ promise further surprises from this puzzling Hsp70 chaperone system.

## Data availability

EM maps have been deposited in the Electron Microscopy Data Bank under accession codes EMDB: EMD-14479 for RAC conformation 1 and EMDB: EMD-14480 for RAC conformation 2. The atomic models have been deposited in the Protein Data Bank under accession numbers PDB: 7Z3N and PDB: 7Z3O.

## Acknowledgements

We thank S. Adrian for growing *C. thermophilum* cultures, A. Hendricks for technical support, and S. Pfeffer and members of the Sinning lab for stimulating discussions. Initial cryo-EM data were collected at the ESRF CM01 with support of Dr. Daouda A. K. Traore. Cryo-EM data used for 80S-RAC structure determination were collected at the University of Heidelberg (HDcryoNet) with support from D. Flemming and G. Hofhaus. We acknowledge the data storage service SDS@hd and bwHPC supported by the Ministry of Science, Research and the Arts Baden Württemberg (MWK) and the Deutsche Forschungsgemeinschaft (DFG) through grants INST 35/1314-1 FUGG and INST 35/1134-1 FUGG. This work was supported by the DFG through the Leibniz Programme (SI 586/6-1) to I.S.

## Author contributions

N.K. and E.H. provided the materials and the protocols for native complex pull-outs from *Chaetomium thermophilum.* K.L. cloned, expressed and purified pull-out *Ct*80S-RAC. M.K. purified *Ct*80S ribosomes. G.V.G. optimized sample for cryo-EM preparation. M.K. prepared cryo-EM samples, collected and processed EM data. M.K. and K.W. built the structural models. M.K., K.W., and I.S. interpreted the data. M.K., K.W., and I.S. wrote the manuscript with contributions from all authors. M.K., K.L., K.W., and I.S. planned the study and designed the experiments.

## Materials & Correspondence

Requests should be addressed to.

## METHODS

### Construct design, cloning and expression

The pRSF-duet-ctSSZ-FTpA was used for ectopic integration and expression of SSZ-FTpA in *Chaetomium thermophilum.* SSZ promoter region (628 bases) and open reading frame were amplified by PCR from *Chaetomium thermophilum* genomic DNA and fused to the Flag-TEV-protA tag resulting into the pRSFduet-ctSSZ-FTpA plasmid. *Chaetomium thermophilum* wildtype strain was transformed with the pRSFduet-ctSSZ-FTpA plasmid as described^25^. In brief, protoplast were generated from the cell wall digestion of the fungus mycelium and mixed with the linearized plasmid DNA. The transformed protoplast were plated and selected on CCM-sorbitol agar plates, supplemented with 0.5mg/ml terbinafine, incubated at 50°C for three days. Expression of the SSZ-FTpA protein was verified by Western blotting of whole-cell lysate using PAP (Sigma-Aldrich, P1291) antibodies according to the manufacturer’s protocol.

ctSSZ1-FTpA *mycelia* were cultivated in a rotary shaker at 90 r.p.m. at 55 °C, harvested through a metal sieve, washed with water, dried with a vacuum filter and immediately frozen in liquid nitrogen. Frozen mycelium cells were ground to fine powder by Cryo Mill (Retch) (5 min, frequency 30/s) and stored at −80 °C.

### Purification of *C. thermophilum* 80S-RAC complexes

The powdered mycelium was resuspended in 20 mM HEPES-KOH (pH 8.0), 150 mM NaCl, 50 mM KOAc, 2 mM Mg(OAc)_2_, 1 mM DTT, 5% glycerol and 0.1% NP-40. Insoluble material was removed by centrifugation (17,000 r.p.m., JA25-50 rotor (Beckman), 30 min). The lysate was transferred onto IgG beads and incubated at 4 °C, for 15 hours. Beads were washed (20 mM HEPES-KOH (pH 8.0), 150 mM NaCl, 50 mM KOAc, 2 mM Mg(OAc)_2_, 1 mM DTT, 5% glycerol, 0.01% NP-40), incubated with TEV protease at 4 °C, for 4 hours and eluted. The elution fractions were pooled together and precipitated by adding 7% w/v of PEG20000. After a 10 min centrifugation, the pellets were resuspended in 20 mM HEPES-KOH (pH 7.5), 50 mM KOAc, 5 mM Mg(OAc)_2_, 2 mM DTT and used for cryo-EM grid preparation or stored at −80 °C.

### Purification of *C. thermophilum* 80S ribosomes

The protocol for the isolation of *Ct*80S ribosomes was as previously described^26^. In brief, the powdered mycelium was resuspended in 20 mM HEPES-KOH (pH 7.5), 500 mM KOAc, 5 mM Mg(OAc)_2_, 2 mM DTT and 0.5 mM PMSF and vortexed until no clumps remained. Insoluble material was removed by centrifugation (20,000 r.p.m., JA25-50 rotor (Beckman), 35 min). Ribosomes were pelleted through a high-salt sucrose cushion (20 mM HEPES-KOH (pH 7.5), 500 mM KOAc, 1.5 M sucrose, 5 mM Mg(OAc)_2_ and 2 mM DTT) at 35,000 r.p.m. in a Ti-865 rotor (Thermo Scientific) for 18 h before they were resuspended in 20 mM HEPES-KOH (pH 7.5), 50 mM KOAc, 5 mM Mg(OAc)_2_, 2 mM DTT and 0.5 mM PMSF. The *Ct*80S ribosomes were then incubated with 1 mM neutralized puromycin solution and 1 mM GTP for 1 hour at 30 °C. The solution containing ribosomes were further purified in 15–40% sucrose gradient (20 mM HEPES-KOH (pH 7.5), 150 mM KOAc, 5 mM Mg(OAc)_2_, 15–40% sucrose, 2 mM DTT and 0.5 mM PMSF) at 18,000 r.p.m. in a Superspin 630 rotor (Sorvall) for 15 h. Peak fractions containing *Ct*80S were pooled together and precipitated by adding 7% w/v of PEG20000. After a 10 min centrifugation, the pellets were resuspended in 20 mM HEPES-KOH (pH 7.5), 50 mM KOAc, 5 mM Mg(OAc)_2_, 2 mM DTT and 0.5 mM PMSF, and stored at −80 °C.

### Cryo-electron microscopy grid preparation and data collection

Three microliters of *C*t80S-RAC pull-out sample at 200 nM concentration was applied on holey carbon grids (Quantifoil R2/1 grid, Quantifoil Micro Tools, GmbH) and plunged-frozen into liquid ethane using a Vitrobot (FEI). The Vitrobot environment chamber was programmed to maintain a temperature of 4 °C and 90% humidity. Initial cryo-EM data were collected at the ESRF CM01 and was used for sample optimization and grid improvement. Cryo-EM data used for the determination of the structures of *Ct*80S-RAC were collected on an in-house Titan Krios (FEI) operating at 300 kV. Data were collected on a Quantum-K3 detector using counting mode. The images were acquired at a nominal magnification of x81,000, with a total dose of 20.6 e^-^/Å^2^. Defocus range was set from −0.8 to −2.5 and every movie was fractioned into 149 frames.

### Single particle analysis and model building

A total of 6,662 micrographs were used for the *Ct*80S-RAC structure determination. The frames were aligned and summed using MotionCor2 whole-image motion correction software^42^. CTFFIND4 was used for contrast transfer function (CTF) estimation of unweighted micrographs^43^. Particle auto-picking was performed with Relion 3.1^44^ (Laplacian-of-Gaussian detection) and inspected manually where majority miss-picked particles or contaminants were removed. Later, particles were extracted (480×480 pixels), down-sampled (120×120 pixels) and subjected to two rounds of reference-free 2D classification in Relion 3.1. First cycle of 2D classification was performed with large search range (20 pixels) to achieve the best possible centering of the particles. The second round was performed in higher precision on 2 times down-sampled particles (240×240 pixels) with smaller search ranges (5 pixels). Only properly centered class averages were selected for subsequent processing steps. Further processing was performed with cisTEM^45^. The stack of 837,930 particles from 2D classification was imported to cisTEM and auto-refined using a yeast 80S ribosome as a reference (low pass filtered to 30 Å). Auto-refined particles were subjected to 3D classification, which resulted in removal of 19% of particles that did not contain RAC. Remaining particles were subjected to additional rounds of 3D focus classification (focusing on RAC or 40S subunit). The final resolution was measured by FSC at 0.143 value as implemented in cisTEM. The local resolution variations were calculated with ResMap^46^. The 80S ribosome model was refined from the *Ct*80S structure (PDB ID: 7OLC)^26^. As starting models for *Ct*RAC building we used the crystal structures of Ssz1 (PDB ID: 6SR6)^8^ and the components of Zuo1 (PDB IDs: 6SR6^8^, 5DJE^22^, 4GMQ^20^, 2LWX^27^). The models were manually built and corrected in Coot^47^, and the real-space refinement was used in Phenix^48^. Atomic models were validated using Phenix and MolProbity^49^.

### Figure preparation

Figures were prepared in GraphPad Prism, Pymol, UCSF Chimera^50^ and UCSF ChimeraX^51^.

## REFERENCES

1. Kramer, G., Shiber, A. & Bukau, B. Mechanisms of Cotranslational Maturation of Newly Synthesized Proteins. Annu Rev Biochem 88, 337–364 (2019).

2. Balchin, D., Hayer-Hartl, M. & Hartl, F.U. In vivo aspects of protein folding and quality control. Science 353, aac4354 (2016).

3. Zhang, Y., Sinning, I. & Rospert, S. Two chaperones locked in an embrace: structure and function of the ribosome-associated complex RAC. Nat Struct Mol Biol 24, 611–619 (2017).

4. Gautschi, M. et al. RAC, a stable ribosome-associated complex in yeast formed by the DnaK-DnaJ homologs Ssz1p and zuotin. Proc Natl Acad Sci U S A 98, 3762–7 (2001).

5. Huang, P., Gautschi, M., Walter, W., Rospert, S. & Craig, E.A. The Hsp70 Ssz1 modulates the function of the ribosome-associated J-protein Zuo1. Nat Struct Mol Biol 12, 497–504 (2005).

6. Weyer, F.A., Gumiero, A., Gese, G.V., Lapouge, K. & Sinning, I. Structural insights into a unique Hsp70-Hsp40 interaction in the eukaryotic ribosome-associated complex. Nat Struct Mol Biol 24, 144–151 (2017).

7. Mayer, M.P. Hsp70 chaperone dynamics and molecular mechanism. Trends Biochem Sci 38, 507–14 (2013).

8. Zhang, Y. et al. The ribosome-associated complex RAC serves in a relay that directs nascent chains to Ssb. Nat Commun 11, 1504 (2020).

9. Yan, W. et al. Zuotin, a ribosome-associated DnaJ molecular chaperone. EMBO J 17, 4809–17 (1998).

10. Cyr, D.M., Langer, T. & Douglas, M.G. DnaJ-like proteins: molecular chaperones and specific regulators of Hsp70. Trends Biochem Sci 19, 176–81 (1994).

11. Cheetham, M.E. & Caplan, A.J. Structure, function and evolution of DnaJ: conservation and adaptation of chaperone function. Cell Stress Chaperones 3, 28–36 (1998).

12. Craig, E.A. & Marszalek, J. How Do J-Proteins Get Hsp70 to Do So Many Different Things? Trends Biochem Sci 42, 355–368 (2017).

13. Rosenzweig, R., Nillegoda, N.B., Mayer, M.P. & Bukau, B. The Hsp70 chaperone network. Nat Rev Mol Cell Biol 20, 665–680 (2019).

14. Kampinga, H.H. & Craig, E.A. The HSP70 chaperone machinery: J proteins as drivers of functional specificity. Nat Rev Mol Cell Biol 11, 579–92 (2010).

15. Gautschi, M., Mun, A., Ross, S. & Rospert, S. A functional chaperone triad on the yeast ribosome. Proc Natl Acad Sci U S A 99, 4209–14 (2002).

16. Hundley, H. et al. The in vivo function of the ribosome-associated Hsp70, Ssz1, does not require its putative peptide-binding domain. Proc Natl Acad Sci U S A 99, 4203–8 (2002).

17. Doring, K. et al. Profiling Ssb-Nascent Chain Interactions Reveals Principles of Hsp70-Assisted Folding. Cell 170, 298–311 e20 (2017).

18. Willmund, F. et al. The cotranslational function of ribosome-associated Hsp70 in eukaryotic protein homeostasis. Cell 152, 196–209 (2013).

19. Peisker, K. et al. Ribosome-associated complex binds to ribosomes in close proximity of Rpl31 at the exit of the polypeptide tunnel in yeast. Mol Biol Cell 19, 5279–88 (2008).

20. Leidig, C. et al. Structural characterization of a eukaryotic chaperone--the ribosome-associated complex. Nat Struct Mol Biol 20, 23–8 (2013).

21. Zhang, Y. et al. Structural basis for interaction of a cotranslational chaperone with the eukaryotic ribosome. Nat Struct Mol Biol 21, 1042–6 (2014).

22. Lee, K., Sharma, R., Shrestha, O.K., Bingman, C.A. & Craig, E.A. Dual interaction of the Hsp70 J-protein cochaperone Zuotin with the 40S and 60S ribosomal subunits. Nat Struct Mol Biol 23, 1003–1010 (2016).

23. Shrestha, O.K. et al. Structure and evolution of the 4-helix bundle domain of Zuotin, a J-domain protein co-chaperone of Hsp70. PLoS One 14, e0217098 (2019).

24. Lee, K. et al. Pathway of Hsp70 interactions at the ribosome. Nat Commun 12, 5666 (2021).

25. Kellner, N. et al. Developing genetic tools to exploit Chaetomium thermophilum for biochemical analyses of eukaryotic macromolecular assemblies. Sci Rep 6, 20937 (2016).

26. Kisonaite, M., Wild, K., Lapouge, K., Ruppert, T. & Sinning, I. High-resolution structures of a thermophilic eukaryotic 80S ribosome reveal atomistic details of translocation. Nat Commun 13, 476 (2022).

27. Ducett, J.K. et al. Unfolding of the C-terminal domain of the J-protein Zuo1 releases autoinhibition and activates Pdr1-dependent transcription. J Mol Biol 425, 19–31 (2013).

28. Mayer, M.P. & Gierasch, L.M. Recent advances in the structural and mechanistic aspects of Hsp70 molecular chaperones. J Biol Chem 294, 2085–2097 (2019).

29. Kityk, R., Kopp, J. & Mayer, M.P. Molecular Mechanism of J-Domain-Triggered ATP Hydrolysis by Hsp70 Chaperones. Mol Cell 69, 227–237 e4 (2018).

30. Zhang, W., Dunkle, J.A. & Cate, J.H. Structures of the ribosome in intermediate states of ratcheting. Science 325, 1014–7 (2009).

31. Punjani, A. & Fleet, D.J. 3D variability analysis: Resolving continuous flexibility and discrete heterogeneity from single particle cryo-EM. J Struct Biol 213, 107702 (2021).

32. Grotwinkel, J.T., Wild, K., Segnitz, B. & Sinning, I. SRP RNA remodeling by SRP68 explains its role in protein translocation. Science 344, 101–4 (2014).

33. Kaschner, L.A., Sharma, R., Shrestha, O.K., Meyer, A.E. & Craig, E.A. A conserved domain important for association of eukaryotic J-protein co-chaperones Jjj1 and Zuo1 with the ribosome. Biochim Biophys Acta 1853, 1035–45 (2015).

34. Gumiero, A. et al. Interaction of the cotranslational Hsp70 Ssb with ribosomal proteins and rRNA depends on its lid domain. Nat Commun 7, 13563 (2016).

35. Kityk, R., Kopp, J., Sinning, I. & Mayer, M.P. Structure and dynamics of the ATP-bound open conformation of Hsp70 chaperones. Mol Cell 48, 863–74 (2012).

36. Hanebuth, M.A. et al. Multivalent contacts of the Hsp70 Ssb contribute to its architecture on ribosomes and nascent chain interaction. Nat Commun 7, 13695 (2016).

37. Jiang, J. et al. Structural basis of J cochaperone binding and regulation of Hsp70. Mol Cell 28, 422–33 (2007).

38. Qi, R. et al. Allosteric opening of the polypeptide-binding site when an Hsp70 binds ATP. Nat Struct Mol Biol 20, 900–7 (2013).

39. Liu, Q. & Hendrickson, W.A. Insights into Hsp70 chaperone activity from a crystal structure of the yeast Hsp110 Sse1. Cell 131, 106–20 (2007).

40. Rakwalska, M. & Rospert, S. The ribosome-bound chaperones RAC and Ssb1/2p are required for accurate translation in Saccharomyces cerevisiae. Mol Cell Biol 24, 918–697 (2004).

41. Chernoff, Y.O. & Kiktev, D.A. Dual role of ribosome-associated chaperones in prion formation and propagation. Curr Genet 62, 677–685 (2016).

## REFERENCES

42. Zheng, S.Q. et al. MotionCor2: anisotropic correction of beam-induced motion for improved cryo-electron microscopy. Nat Methods 14, 331–332 (2017).

43. Rohou, A. & Grigorieff, N. CTFFIND4: Fast and accurate defocus estimation from electron micrographs. J Struct Biol 192, 216–21 (2015).

44. Scheres, S.H. RELION: implementation of a Bayesian approach to cryo-EM structure determination. J Struct Biol 180, 519–30 (2012).

45. Grant, T., Rohou, A. & Grigorieff, N. cisTEM, user-friendly software for single-particle image processing. Elife 7 (2018).

46. Kucukelbir, A., Sigworth, F.J. & Tagare, H.D. Quantifying the local resolution of cryo-EM density maps. Nat Methods 11, 63–5 (2014).

47. Emsley, P., Lohkamp, B., Scott, W.G. & Cowtan, K. Features and development of Coot. Acta Crystallogr D Biol Crystallogr 66, 486–501 (2010).

48. Adams, P.D. et al. PHENIX: a comprehensive Python-based system for macromolecular structure solution. Acta Crystallogr D Biol Crystallogr 66, 213–21 (2010).

49. Williams, C.J. et al. MolProbity: More and better reference data for improved all-atom structure validation. Protein Sci 27, 293–315 (2018).

50. Pettersen, E.F. et al. UCSF Chimera--a visualization system for exploratory research and analysis. J Comput Chem 25, 1605–12 (2004).

51. Pettersen, E.F. et al. UCSF ChimeraX: Structure visualization for researchers, educators, and developers. Protein Sci 30, 70–82 (2021).

