## Supplemental Data for "Structural inventory of cotranslational protein folding by the eukaryotic RAC complex"

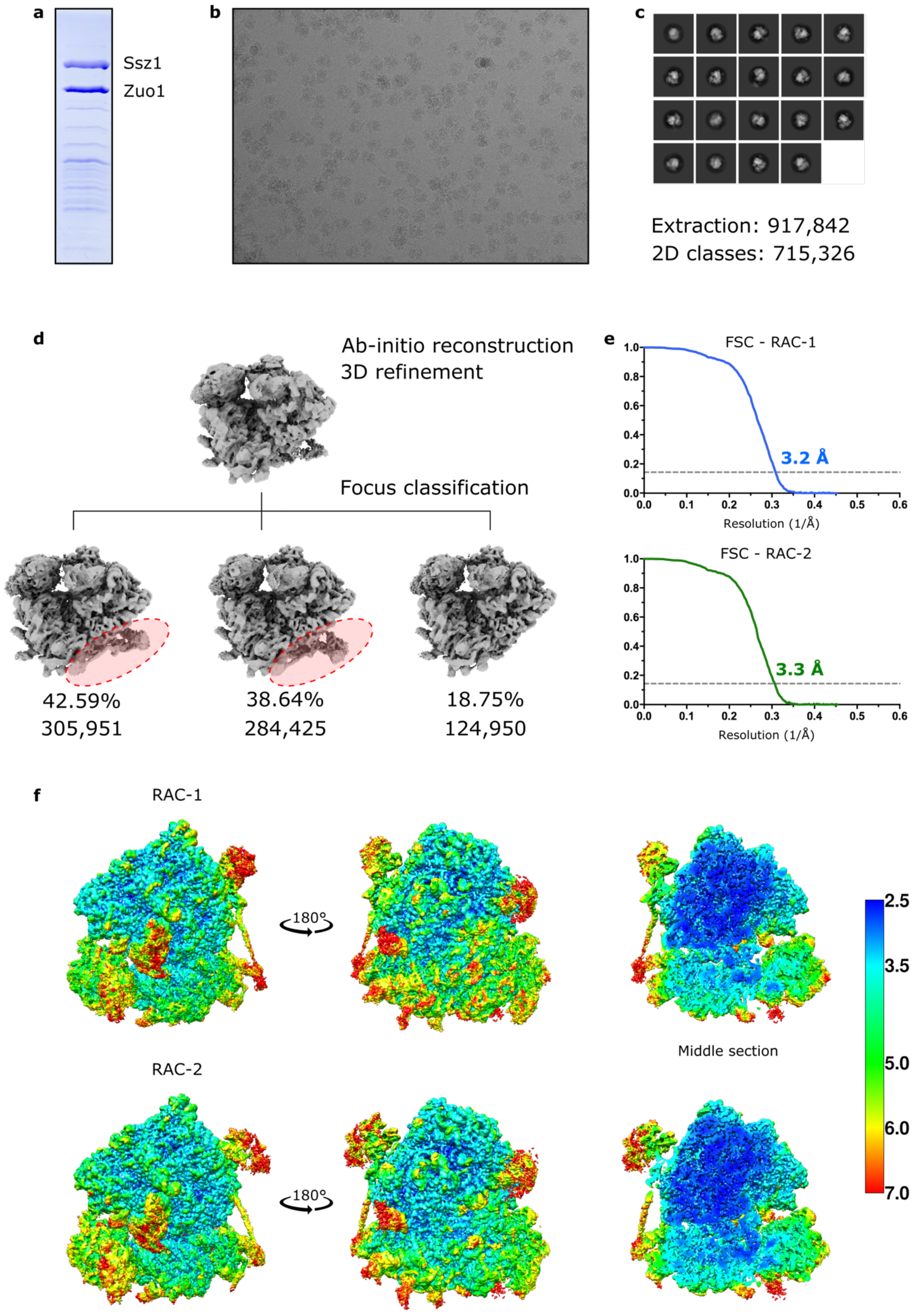
a
Ssz1
Zuo1
b
c
Extraction: 917,842
2D classes: 715,326
d
Ab-initio reconstruction
3D refinement
Focus classification
42.59%
305,951
38.64%
284,425
18.75%
124,950
e
FSC - RAC-1
3.2 Å
Resolution (1/Å)
FSC - RAC-2
3.3 Å
Resolution (1/Å)
f
RAC-1
180°
Middle section
RAC-2
180°
Middle section
2.5
3.5
5.0
6.0
7.0

**Extended Data Figure 1 | Sample preparation of *C. thermophilum* RAC-80S and processing of the cryo-EM data.**

**a,** Purified sample from *C. thermophilum* Ssz1 pull-out analyzed by SDS-PAGE and Coomassie staining. The analysis shows prominent bands of Ssz1, Zuo1 and also 80S ribosomal proteins. **b,** Cryo-EM micrograph of the RAC-80S ribosome complexes. **c-d,** 2D classes and a flow chart showing the stages of cryo-EM image processing. A total of 8,432 micrographs was collected on a Titan Krios 300 kV microscope and subjected to beam-induced motion correction. A total of 917,842 particles were auto-picked and after multiple rounds of 2D classification resulted in a selection of 715,326 particles. The particles were subjected to 3D refinement and 3D classification focused on RAC. Data processing was done using RELION 3.0 and cryoSPARC (v3.2). **e,** The 0.143 Fourier Shell Correlation (FSC) cut-off criteria indicates that the cryo-EM maps of the RAC-80S ribosome in RAC-1 and RAC-2 conformations have average resolutions of 3.2 Å and 3.3 Å, respectively. **f,** Local resolution distribution displayed on the reconstructed cryo-EM density maps of the RAC-1 and RAC-2 complexes.

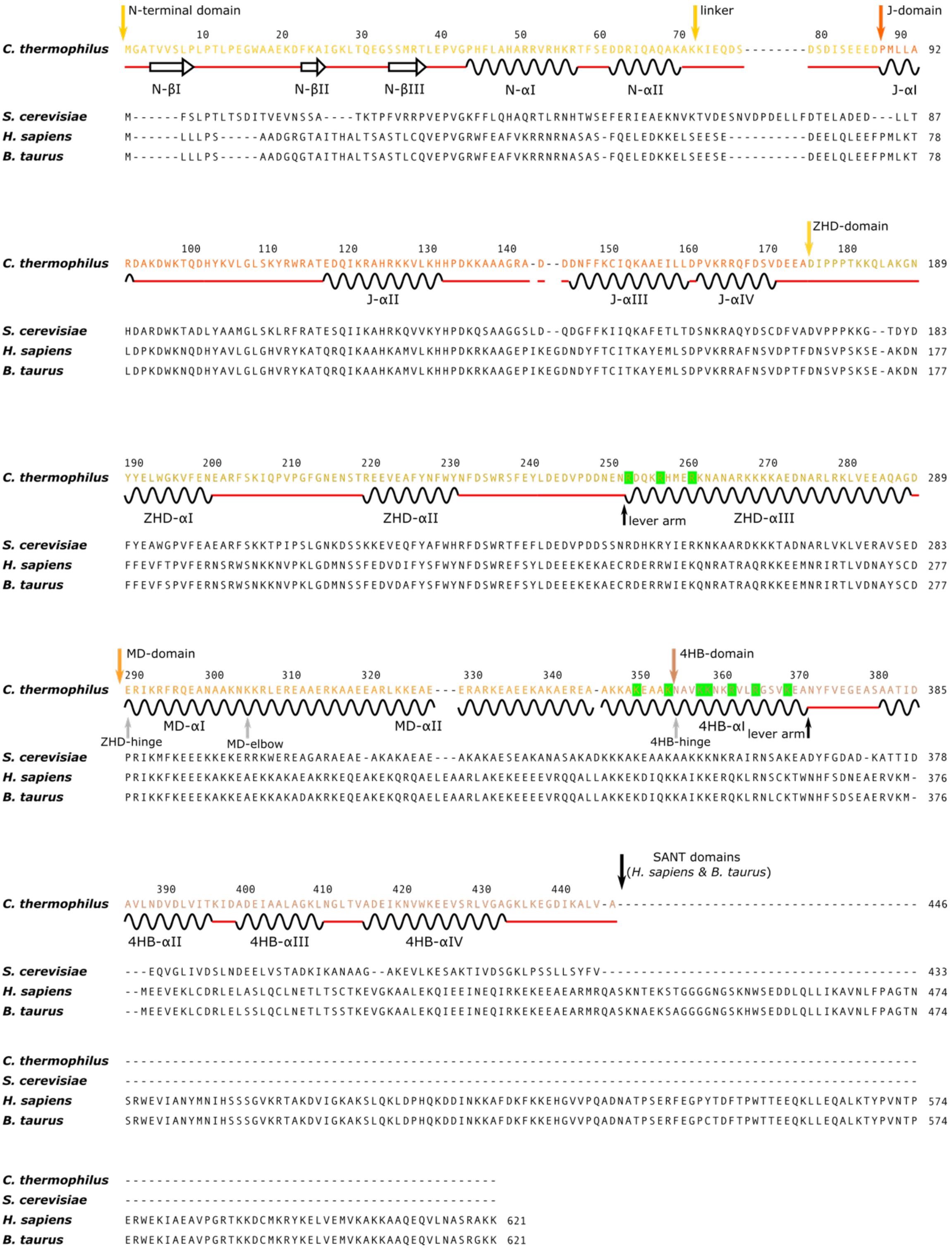


**Extended Data Figure 2 | Multiple sequence alignment and secondary structure of *C. thermophilum* Zuo1.** Sequence alignment of *C. thermophilum* Zuo1 with *S. cerevisiae*, *H. sapiens* and *B. taurus* homologs. Domains are marked with arrows above the sequence and colored accordingly. Specific kinks in the Zuo1 lever arm are indicated by grey arrows. Residues colored in green are important for two anchoring interactions of Zuo1 to the 80S ribosome.

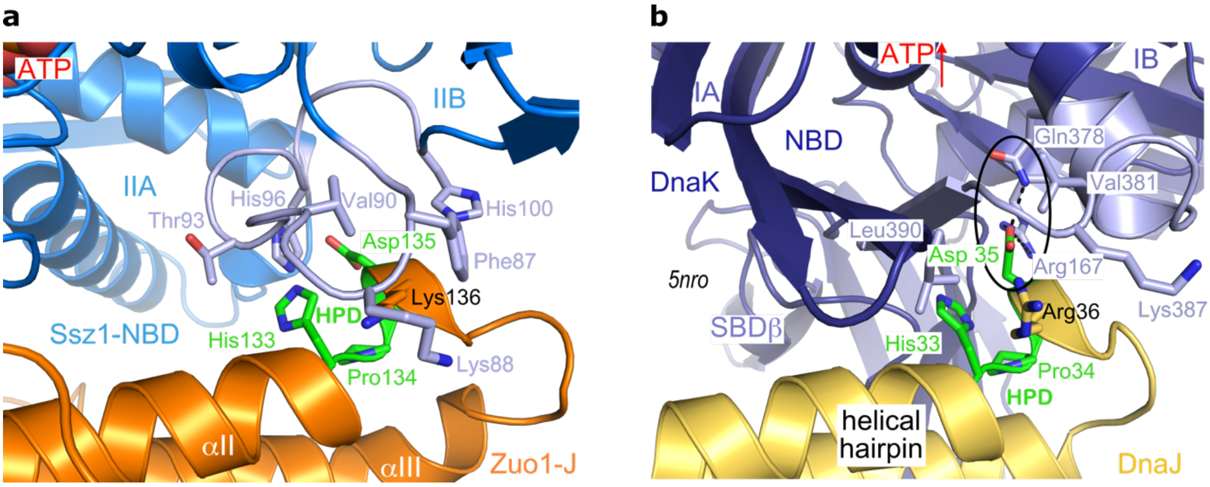


**Extended Data Figure 3 | The HPD motif within the Hsp40/Hsp70 interaction**
The conserved HPD motif within a helical hairpin of Hsp40 J-domains forms the central element for the activation of ATP-hydrolysis in Hsp70 proteins. **a,** Zoom into the Ssz1-NBD/Zuo1-J interface of the RAC-1 complex (same in RAC-2). Ssz-1 binds to but does not hydrolyze ATP. Residues involved in the interface around the HPD motif (green) are detailed. **b,** Zoom into the activating DnaK/DnaJ interface[1], which is different and includes the linker of DnaK-NBD to its SBDβ. The central interaction network of the aspartate within the HPD motif is encircled. The position of ATP is indicated by an arrow.

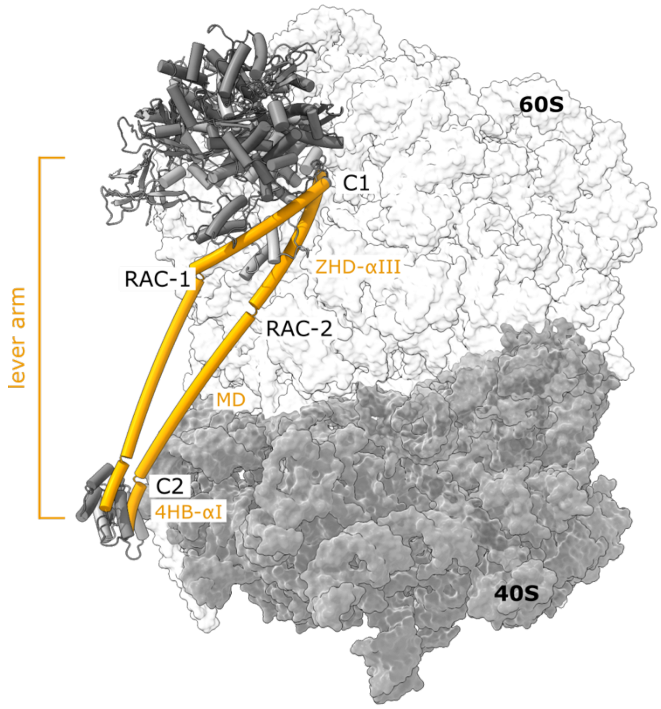


**Extended Data Figure 4 | Two RAC conformations on the 80S ribosome.**
RAC adopts two discrete conformations (RAC-1 and RAC-2) on the 80S ribosome, shown here in superposition. Interactions with the ribosome are formed through multiple contacts of a Zuo1 lever arm (shown in orange cartoon) that includes the helix αIII of the ZHD, the entire MD, and the helix αI of the 4HB. Ribosome is shown in surface representation.

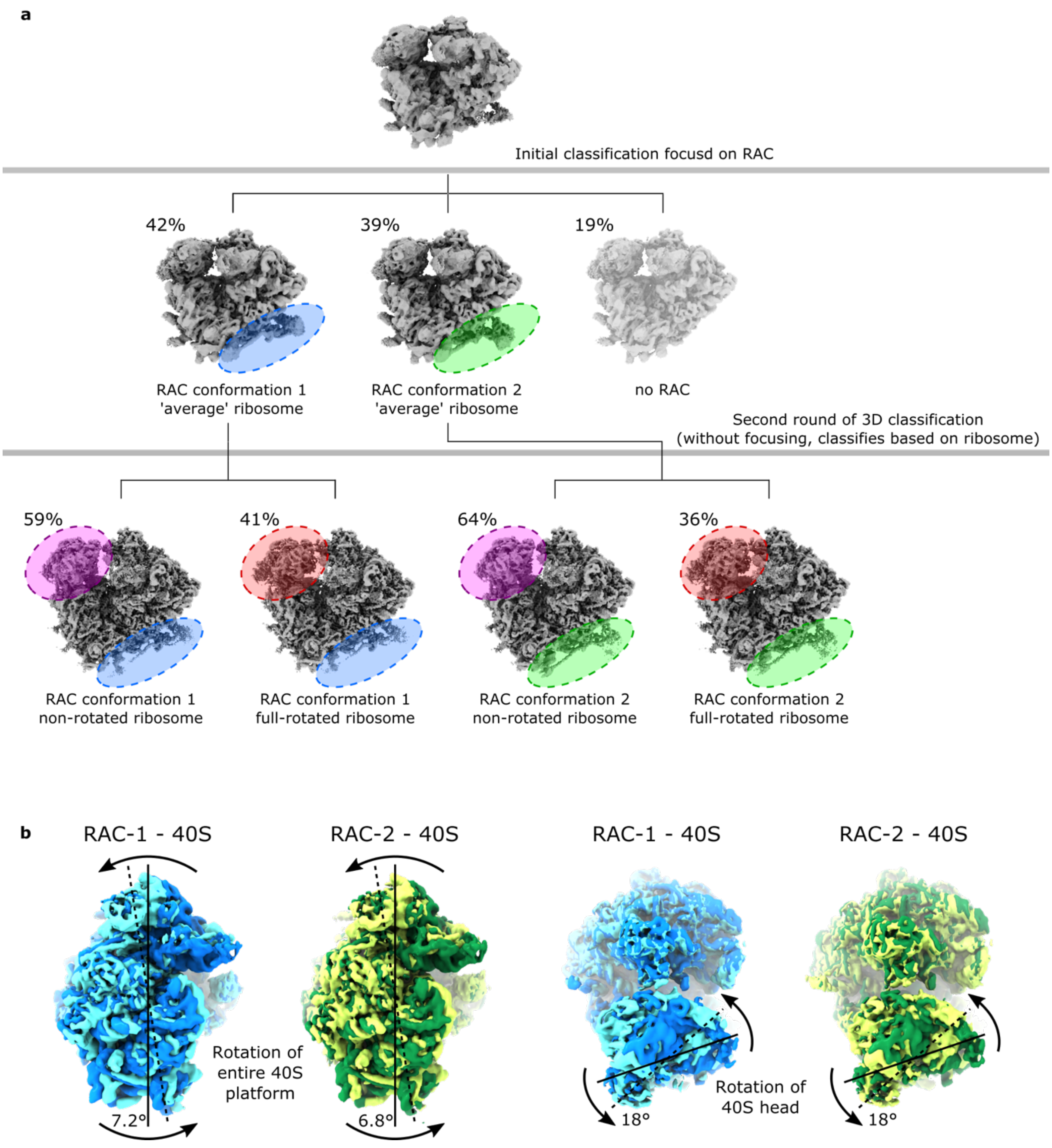


**Extended Data Figure 5 | Extended processing of the cryo-EM data and 3D variability analysis.**

**a,** Two 3D classes containing RAC (see Extended Data Fig. 1, here marked with blue and green circles) were further analyzed by a second round of 3D classification focused on the ribosome 40S head (marked with red and magenta circles). Such classification allowed to separate fully rotated 80S ribosomes from non-rotated 80S particles (40S head is main indication in the rotation movement). Classification showed that both RAC conformations (RAC-1 and RAC-2) can accommodate full ribosomal rotation, and contain similar percentage of full-rotated and non-rotated 80S ribosomes. **b,** 3D variability analysis of ribosomal rotation. RAC-1 is shown in blue colors, and RAC-2 is shown in green. Overlaid maps (low-pass filtered) are the first frame and the last frame of the 3D variability analysis. Top panels show the rotation of the 40S body (about 7°), and bottom panels visualize the rotations (swiveling) of the 40S head (when body is aligned, 18°). Rotations were measured in UCSF-Chimera using the command ‘measure rotation‘.

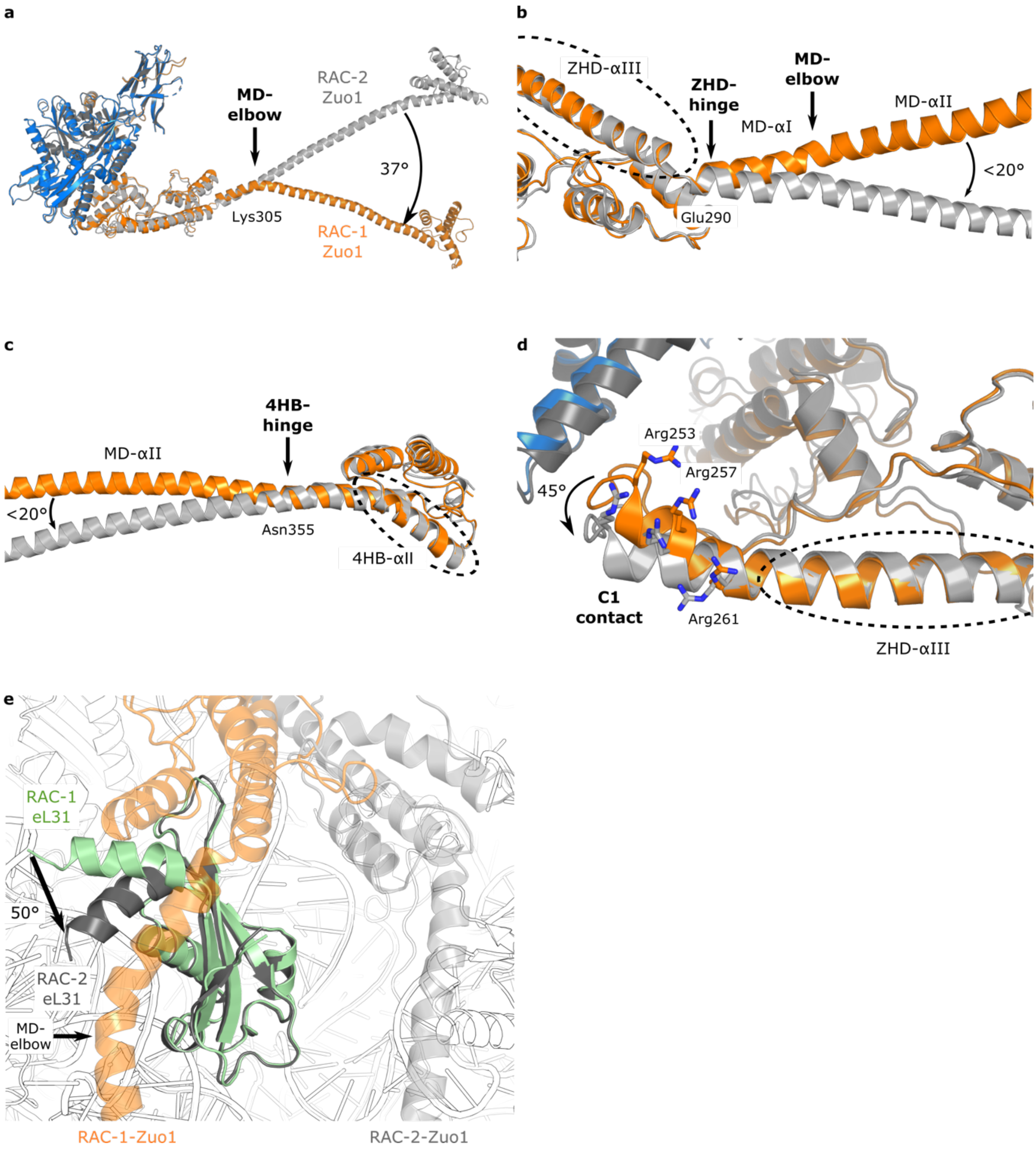


**Extended Data Figure 6 | Rearrangements of the Zuo1 helical lever arm and ZHD anchor region in different RAC conformations. a,** Full view of the *C. thermophilum* RAC, when superposed on Ssz1. RAC-1 corresponds to a bent MD-elbow (orange) and RAC-2 to a straightened arm (gray). **b-c,** Zoom in views of ZHD-hinge, MD-elbow and 4HB-hinge. **d,** Rearrangements in the Zuo1-ZHD anchor region (45° rotation). In **b** to **d** superposition is performed on the areas marked by dashed circle**. e,** Changes in eL31 N-terminal helix in different RAC conformations. The N-terminal helix of eL31 is rotated by 50° in RAC-1 in respect to RAC-2. RAC-1 eL31 is shown in green cartoon, RAC-2 eL31 is given in grey. The ribosome is depicted as a transparent cartoon, with Zuo1 shown in orange.

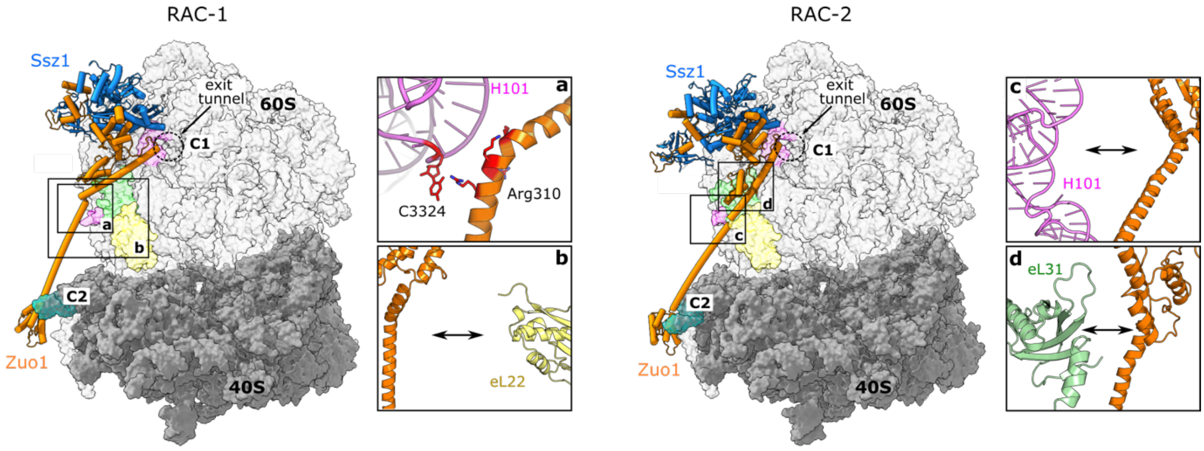


**Extended Data Figure 7 | RAC interactions with the 80S ribosome.**
The main interaction sites of RAC-1 and RAC-2 (shown in the main panels) are highlighted with squares that correspond to the zoom images **a** to **d**. **a, c,** Zuo1 MD-$\alpha$I interaction with H101 of 26S rRNA in RAC-1 (**a**) and its abolishment in RAC-2 (**c**). **b,** Zuo1-ZHD domain is shifted away from eL22 in RAC-1 (compare with Fig. 1e for C2). **d,** ZHD-MD domains shifted away from eL31 in RAC-2 (compare with Fig. 1b for RAC-1).

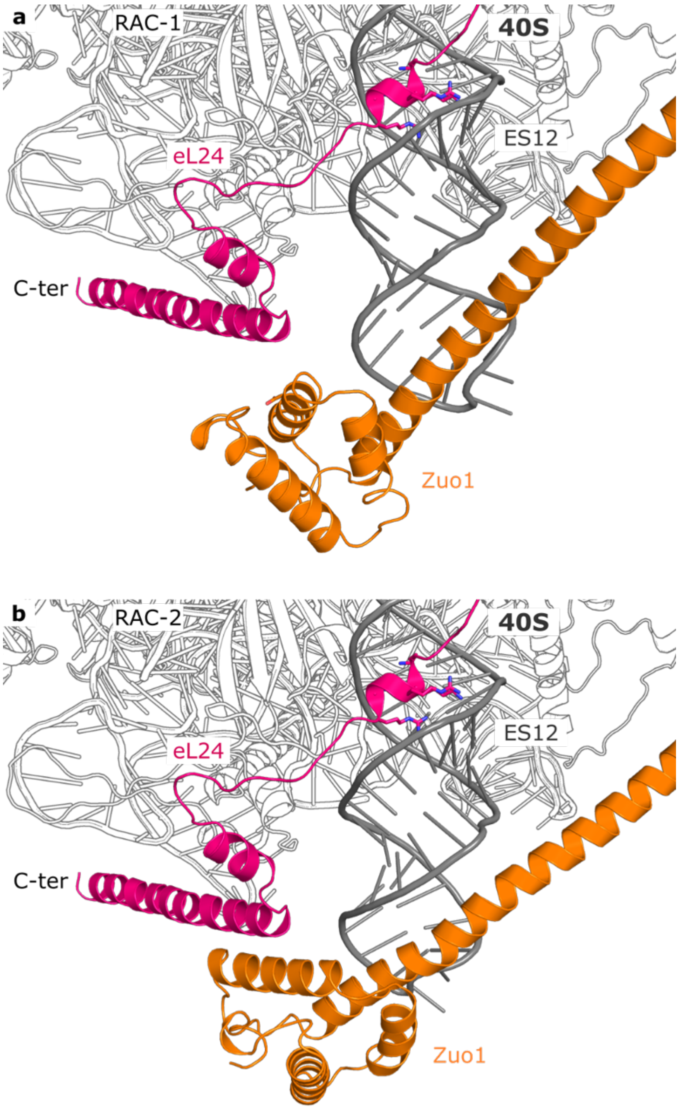


**Extended Data Figure 8 | eL24 interaction with ES12.**

**a-b,** A ribosome-internal arginine-rich motif (ARM) is provided by eL24 of the 60S subunit and holds ES12 in a fixed location by binding in a widened major groove in both RAC-1 (**a**) and RAC-2 (**b**) conformations. The C-terminal helix of the eL24 anchors on the 40S body of the ribosome and is positioned close to the Zuo1-4HB domain.

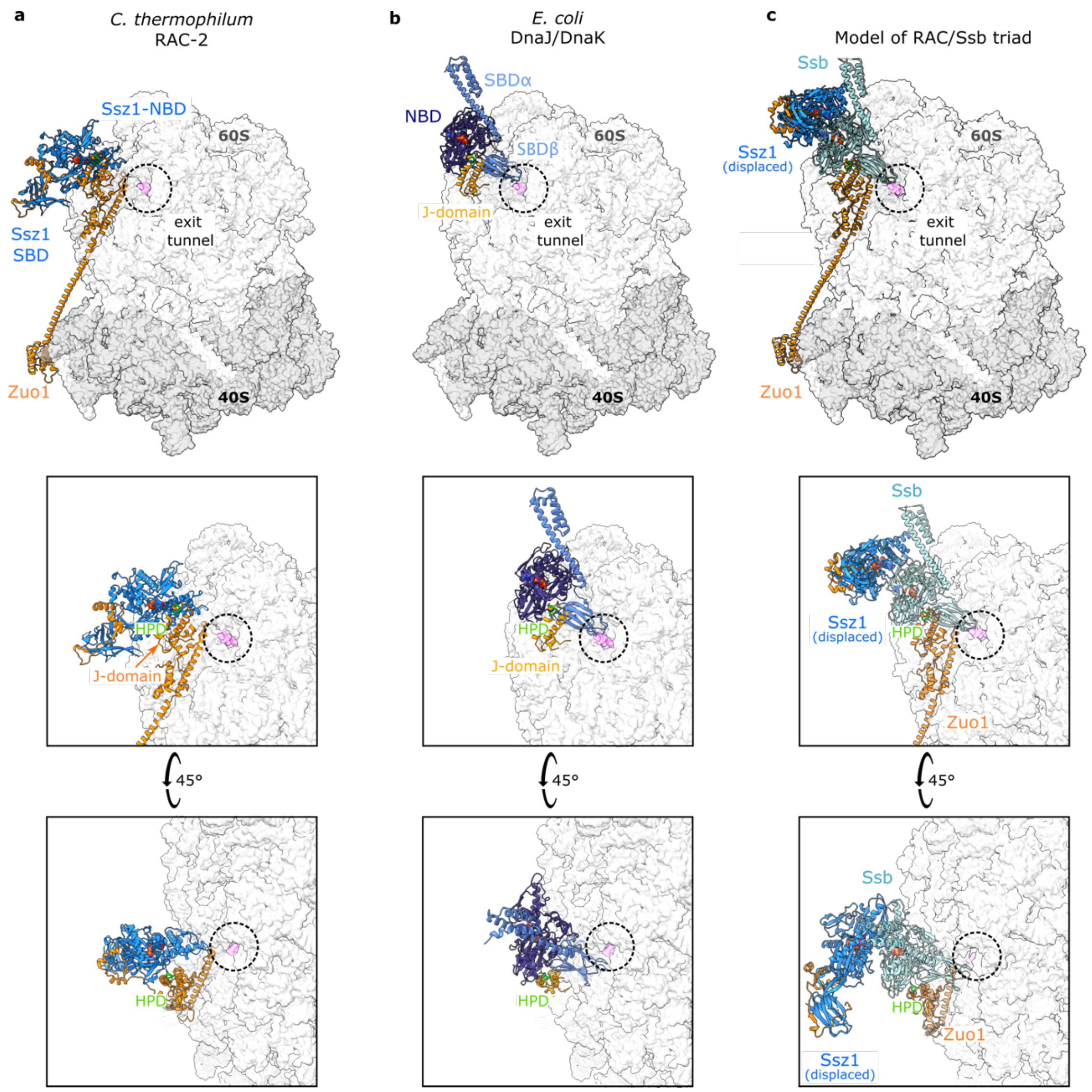


**Extended Data Figure 9 | Model of the RAC/Ssb triad on the ribosome.**
**a,** RAC-80S complex in RAC-2 conformation. The 80S ribosome is shown in a transparent surface representation (60S: pale grey, 40S: dark grey, NC: magenta), Zuo1 is shown as orange cartoon, and Ssz1 as blue cartoon. **b,** The *E. coli* DnaK/DnaJ complex[1], with the J-domain of DnaJ superposed on the Zuo1 J-domain of the RAC-2 complex. **c,** DnaK (from panel **b**) is replaced by Ssb (open, ATP-bound state)[2] to obtain a model for Ssb activation by Zuo1-HPD. In this model, the Ssz1-NBD that masks the Zuo1-HPD motif needs to detach from the Zuo1 J-ZHD at the ribosomal tunnel exit. Thus, Ssz1-NBD is moved to a position matching a crystallographic Ssb homodimer[2] (PDB code 5tky), in agreement with recent Ssz1/Ssb cross-linking studies[3]. Ssz1-SBD bound to Zuo1-N is kept in close neighborhood to the ribosome-docked parts of Zuo1 by the short Zuo1 N-J linker.

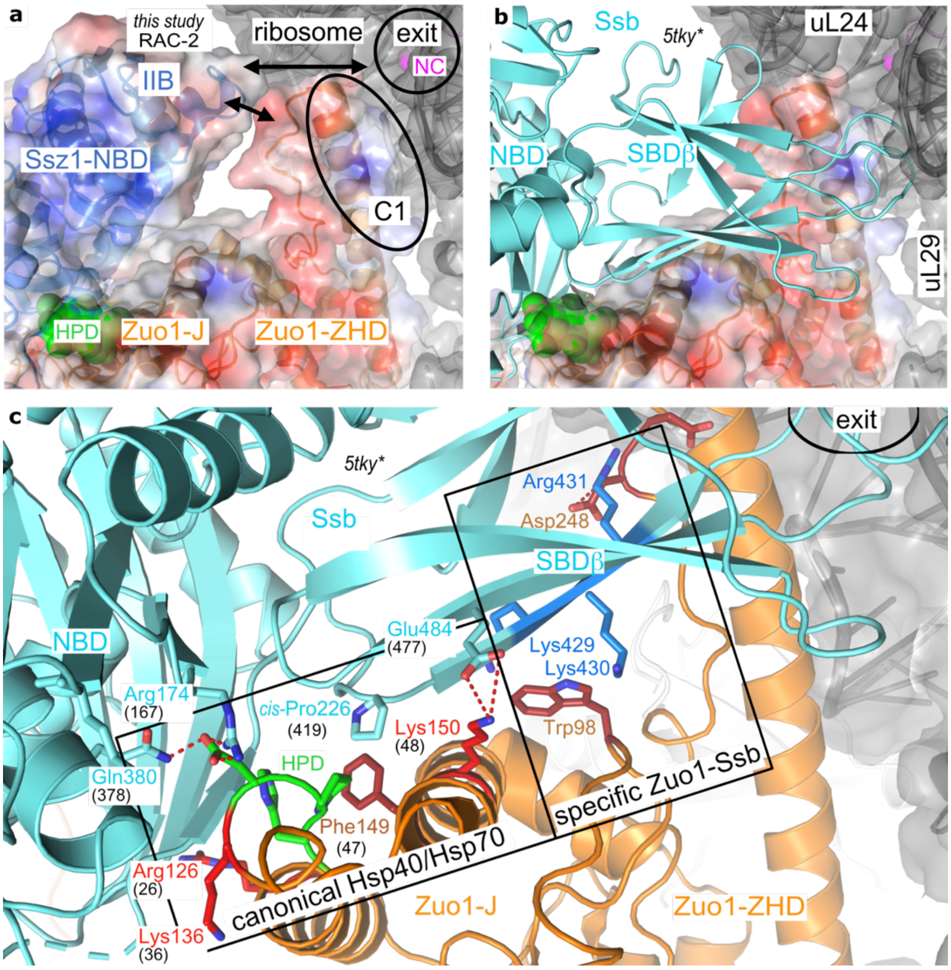


**Extended Data Figure 10 | Model for the activated RAC-Ssb complex.**
**a,** Zoom on the RAC-2 structure at the major C1 contact of Zuo1-ZHD next to the ribosomal tunnel exit. Surfaces are colored for RAC according their surface potentials[4] (±5kT; blue: positive, red: negative). The HPD motif is highlighted in green. In RAC-2, Ssz1-NBD is displaced from the tunnel exit and the contact to Zuo1-ZHD is broken (indicated by arrows). **b,** Same view with Ssb-ATP from *C. thermophilum* (cyan) bound to the HPD motif in the activating position (modelled as described in main text). Ssb-SBDβ locates next to negatively charged surface patches of Zuo1 ready to accept substrate NCs, in line with cross-links between uL24 and uL29 with Ssb-SBDβ[3]. **c,** Zuo1-Ssb interactions extend the canonical Hsp40/Hsp70 contact around the HPD motif by specific interactions of Zuo1-ZHD with Ssb-SBDβ. The interaction in our model involves a conserved basic motif in Ssb-SBDβ (here: KKR 429-431) that has previously been assigned to ribosome binding[5]. Interactions are detailed for the *C. thermophilum* complex (residue numbers in parenthesis correspond to the DnaK-DnaJ complex of *E. coli*[1]).

### Extended Data Table 1. Data, refinement and model statistics

| State | RAC-1 | RAC-2 |
|---|---|---|
| Data collection statistics | | |
| Microscope | Titan Krios | |
| Camera | K3 | |
| Voltage (kV) | 300 | |
| Magnification | 81,000 | |
| Total dose ($e^-$/Å$^2$) | 39.42 | |
| Defocus rage ($\mu$m) | -0.8 to -2.5 | |
| Calibrated pixel size (Å) | 1.1 | |
| Micrographs collected | 8,432 | |
| Initial number of particles | 917,842 | |
| Refined particles | 715,326 | |
| Symmetry | C1 | |
| Particles in final classes | 305,951 | 284,425 |
| Map resolution (Å) | 3.2 | 3.3 |
| Model Composition | | |
| Chains | 88 | 88 |
| Atoms | 213,544 | 213,453 |
| Residues | Protein: 12,785<br>Nucleotide: 5,238 | Protein: 12,787<br>Nucleotide: 5,231 |
| Ligands | ZN: 8 | ZN: 8 |
| | MG: 455 | MG: 472 |
| | ATP: 1 | ATP: 1 |
| Water | 0 | 0 |
| Bonds (RMSD*) | | |
| Length (Å) (# > 4σ) | 0.003 (2) | 0.003 (0) |
| Angles (°) (# > 4σ) | 0.685 (74) | 0.672 (54) |
| MolProbity score | 1.82 | 1.77 |
| Clash score | 10.32 | 8.97 |
| Ramachandran plot (%) | | |
| Favored | 95.88 | 95.84 |
| Allowed | 4.07 | 4.11 |
| Outliers | 0.05 | 0.06 |
| Ramachandran plot Z-score (RMSD) | | |
| Whole | -1.03 (0.07) | -1.04 (0.07) |
| Helix | -0.03 (0.08) | -0.03 (0.08) |
| Sheet | -0.09 (0.13) | -0.20 (0.12) |
| Loop | -1.23 (0.07) | -1.24 (0.07) |
| Rotamer outliers (%) | 0.12 | 0.10 |
| Cβ outliers (%) | 0.02 | 0.00 |
| Peptide plane (%) | | |
| Cis proline/general | 0.0/0.0 | 0.0/0.0 |
| Twisted proline/general | 0.0/0.0 | 0.0/0.0 |
| CaBLAM outliers (%) | 2.81 | 2.77 |
| ADP min/max/mean (Å$^2$) | | |
| Protein | 66.61/710.48/225.32 | 71.11/698.32/185.30 |
| Nucleotide | 68.09/999.99/180.31 | 72.97/795.73/151.46 |
| Ligand | 51.21/451.41/130.84 | 51.78/457.65/121.47 |
| Occupancy (%) | 100 | 100 |
| Model vs. Data (CC mask) | 0.87 | 0.89 |

*RMSD: root-mean-square deviation

**Extended Data Movie 1 | Both RAC conformations on the 80S ribosome can accommodate ribosomal rotation.** The movie is generated by morphing between fully-rotated and non-rotated RAC-80S ribosomes (40S is shown in dark grey cartoon, 60S: pale grey, Zuo1: orange, Ssz1: blue). The movie highlights that both RAC conformations (RAC-1 and RAC-2) can accommodate full ribosomal rotation via the Zuo1-MD domain. The movie was generated using PyMOL software.

### References


1. Kityk, R., Kopp, J. & Mayer, M.P. Molecular Mechanism of J-Domain-Triggered ATP Hydrolysis by Hsp70 Chaperones. *Mol Cell* **69**, 227-237 e4 (2018).
2. Gumiero, A. et al. Interaction of the cotranslational Hsp70 Ssb with ribosomal proteins and rRNA depends on its lid domain. *Nat Commun* **7**, 13563 (2016).
3. Lee, K. et al. Pathway of Hsp70 interactions at the ribosome. *Nat Commun* **12**, 5666 (2021).
4. Jurrus, E. et al. Improvements to the APBS biomolecular solvation software suite. *Protein Sci* **27**, 112-128 (2018)
5. Hanebuth, M.A. et al. Multivalent contacts of the Hsp70 Ssb contribute to its architecture on ribosomes and nascent chain interaction. *Nat Commun* **7**, 13695 (2016).
